# The redesign of the molecular scaffold of viral ion channel blockers

**DOI:** 10.64898/2026.04.30.721843

**Authors:** Balázs Zoltán Zsidó, Erzsébet Mernyák, Fanni Földes, Zoltán Kopasz, Krisztina Leiner, Mónika Madai, Brigitta Zana, Anett Kuczmog, Csaba Hetényi

## Abstract

The rise of new, rapidly mutating viruses presents increasing challenges for drug developers. Traditional methods, such as high-throughput screening and drug repurposing against mutagenic viral targets, have recently shown their limitations. Our current rational molecular engineering approach offers a sustainable solution by targeting viral ion channels, which generally have low mutation rates. First, extending the amantadine molecule led to the development of new compounds that better match the alternating hydrophobic and hydrophilic patterns of the inner walls of ion channels—a common feature across many viruses. Then, simplifying the structure yielded a cyclohexylamine-based minimalist scaffold that effectively blocks the ion channel and demonstrates improved antiviral activity compared to well-known agents such as amantadine and arterolane. SARS-CoV-2 variants served as test systems in laboratory experiments. The new molecular scaffolds presented here provide a strong foundation for designing potent, broad-spectrum viral ion channel blockers.

## Introduction

Antiviral design is hindered by a lack of reliable animal models and a scarcity of non-mutagenic targets [1–3] within the small viral proteome [4]. While enzymes are common targets [5,6], frequent mutations in their binding pockets ([7–9], Fig S1) lead to rapid evolutionary resistance. The low binding affinity [10,11] and narrow spectrum [11,12] of enzyme inhibitors limit their utility as drugs. As seen with nirmatrelvir, even FDA-approved drugs with strong *in vitro* activity [13] can fail to prevent viral reemergence [14], as the virus quickly develops resistance [7,8,15–18] through target mutations [7,8,15–19].

Targeting proteins other than enzymes, particularly ion channels [20,21], offers a strategy to overcome the antiviral challenges [22] outlined above. Viral ion channels are often non-mutagenic (Fig. 1A) or possess only a few mutations that do not affect channel function [23,24]. The internal structure of viral ion channels often features an alternating pattern of hydrophilic and hydrophobic bands (Fig. 2). They can accommodate [25] not only ions but also small drug molecules such as amantadine (AMA) and rimantadine (RIM, Fig. 1B). AMA and RIM were initially approved for the treatment of influenza A [26]. Such ion channel blockers with an **am**phipathic **s**caffold (AMS) are composed of a hydrophilic head (the protonated amino group [26]) and a hydrophobic tail (the bulky adamantyl group [27], Fig. 1B). This chemical match between AMSs and the alternating pattern of ion channel interiors results in an efficient block of ion transport [25,28], and disruption of viral replication [28]. The development of AMS antivirals is promising for at least three main reasons.

**Fig. 1.**
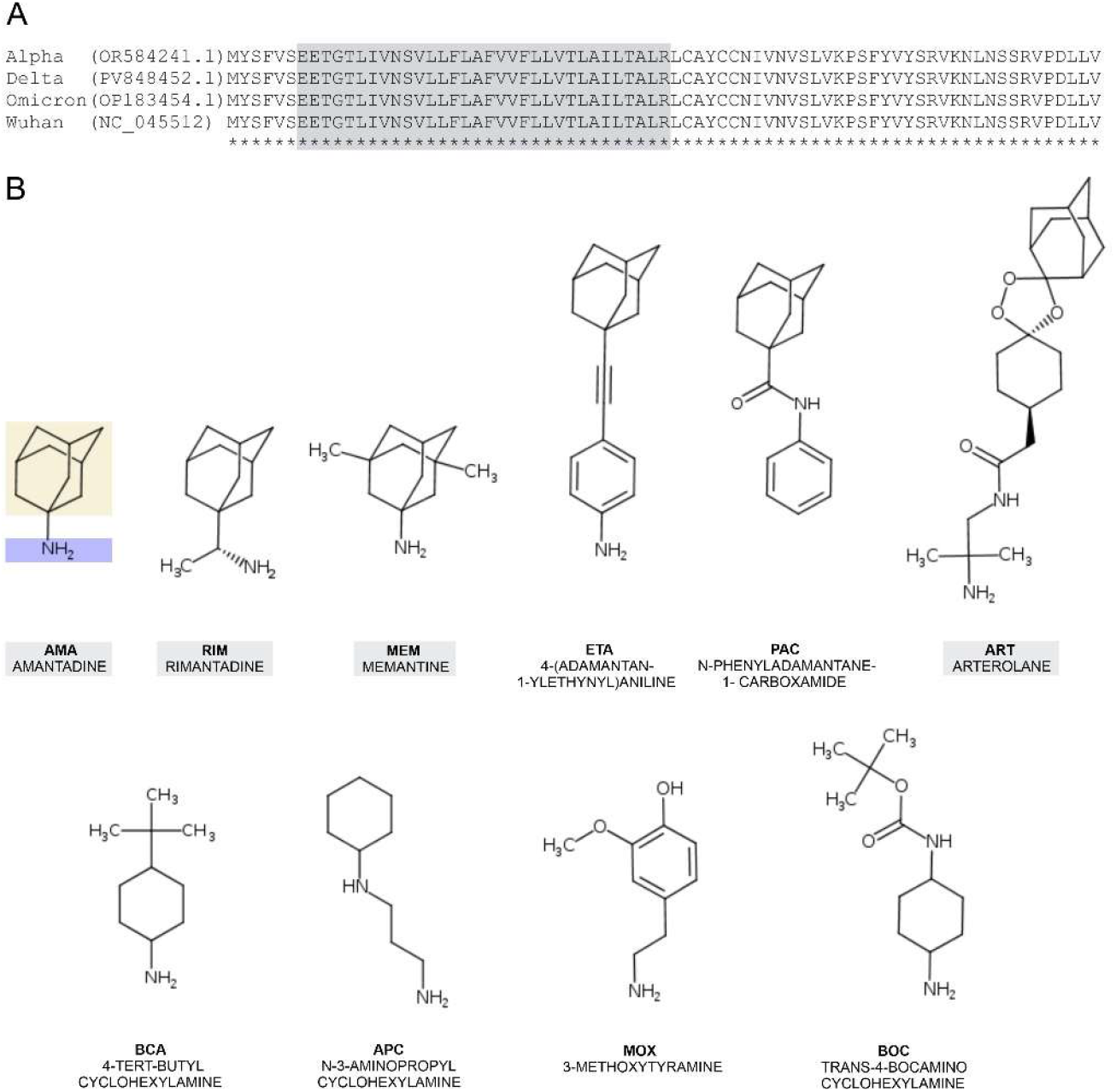
A) The amino acid sequence alignment of the envelope protein (EP), the ion channel of SARS-CoV-2 show no mutations between the different variants. Sequences were obtained from the National Center for Biotechnology Information [29], with repository codes shown in the figure. The transmembrane region is colored grey [28]. Sequence alignment was performed using Clustal Omega [30]. B) The compounds with an **am**phipathic **s**caffold (AMS) investigated in the present study, arranged in increasing order of molecular weight. The compounds in the upper row are based on the AMA scaffold, while those in the lower row have a simplified (minimalist) scaffold. The labels of FDA-approved drugs (used as references) are highlighted in gray. The AMS, composed of a hydrophobic (sandy background) and hydrophilic (blue background) group, is highlighted in AMA.

**Fig. 2.**
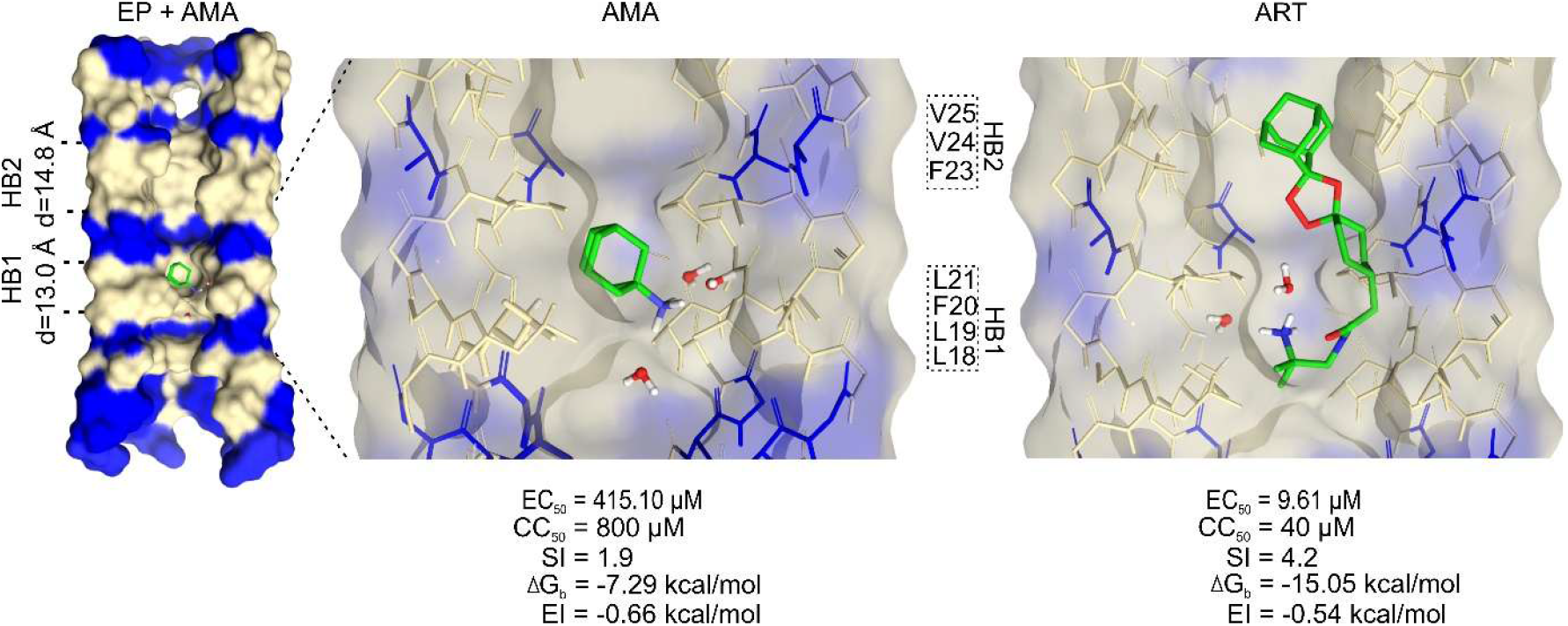
The ion channel (EP) of SARS-CoV-2 binding AMS compounds. Cross-section of EP shown as a surface (left), with AMA (middle), and arterolane (ART) (right) represented with sticks. The blue and tan bands indicate hydrophilic and hydrophobic regions, respectively, based on the chemical properties of the amino acid residues lining the internal wall of EP. The hydrophobic adamantyl group of AMA fits into the hydrophobic bands of EP, while the hydrophilic amino group interacts with the hydrophilic bands via water molecules. These bridging water molecules are represented as red and white balls-and-sticks (not all are shown for clarity). Amino acids are labeled according to PDB entry 7k3g [28]. Hydrophobic bands (HBs) 1 and 2 are labeled, with their diameters shown, and the amino acids highlighted within each HB. The bottom of the figure points towards the extraviral space.

1. Known AMS compounds primarily block non-mutagenic viral ion channels, such as the M2 protein of influenza A [26] and the envelope protein (EP) of SARS-CoV-2 [28,31], which are promising drug targets in the long term. For example, EP is less mutagenic ([24,32–34], Fig. 1A) than other SARS-CoV-2 proteins, such as the spike protein (Fig. S1), which is primarily responsible for vaccine resistance [19].
2. A bulky hydrophobic tail of an AMS can shield nearby functional groups from metabolic cleavage by limiting the access of metabolic enzymes [27,35].
3. The AMS compounds are broad-spectrum antivirals due to their target similarity. They have been effective against various viruses, including influenza [36], SARS-CoV-1 [37], SARS-CoV-2 [25,28,38–42], and hepatitis A [43]. The AMS pattern fits the alternating hydrophilic and hydrophobic bands (Fig. 2) of the inner wall of ion channels present in different viruses [28] allowing broad repurposing.

Given the structural similarity (Point 3) of the targeted ion channels, it was possible to reposition the AMS compounds from M2 of influenza to EP of SARS-CoV-2. This repositioning was first suggested by an NMR study [28] and later supported by *in vitro* and *in vivo* studies [28,38–42] and several clinical trials and clinical case studies [44–50]. The initial insights into how AMA-based compounds bind to EP were provided by NMR [28] and *in silico* [25] studies. Later, several electrophysiology studies verified that the EP is the target of the AMS compounds in SARS-CoV-2 [51–55].

While the examples above show a promising direction, further structure activity relationship (SAR) elucidation is needed (see the author’s remark in ref. [39]) to sustain rational target-based AMS design. Studies with high safety standards at Biological Safety Level 3 (BSL-3, [56]) are necessary to test the antiviral potency of new AMS. Testing of AMS compounds across different variants is completely missing. Furthermore, the synthetic challenges [57] of adamantyl derivatives (Fig. 1B) also hinder the design of new AMS compounds.

Although the AMS-based design appears to be a successful approach, several questions remain unanswered, prompting a re-evaluation of the AMS strategy. The present study addresses these issues and introduces new compounds with an extended, then simplified, synthetically feasible AMS. Their antiviral activity and safety were tested against various SARS-CoV-2 variants in a BSL-4 laboratory.

## Results and Discussion

The rethinking of AMS was carried out through a four-step process in this study. It started with building and validating SARs for known AMS compounds (Step 1). This involved identifying atomic resolution binding modes and calculating their binding affinities. After successfully repositioning M2 to EP, valuable M2-binding and affinity data were also integrated into the SAR development. Experimental test systems were established to verify the virological and cytological activity of the AMS shown in Fig. 1B (Step 2). Using the SARs from Step 1, the AMA scaffold was expanded (Step 3) to better match the ion channel. Finally, simplification (Step 4) resulted in a new, broadly applicable scaffold. In the following Sections, Steps 1-4 will be described in detail.

### 1 Building and validation of SARs

Drug design requires precise SARs based on atomic-resolution structures, but experimental data is currently limited to the influenza M2 channel in complex with AMS compounds [26]. In the case of SARS-CoV-2, only the structure of the unliganded (apo) EP was measured [28] and an *in silico* study [25] explored the binding modes of AMA and RIM to EP using a new protocol, HydroDock, that accounts for interacting water molecules. To calculate the binding mode of other compounds in Fig. 1B, HydroDock [25] was also used in the present study. The quantum mechanics-based QMH-L [58] method was used to calculate binding free energies (ΔG_b_, Table S1). QMH-L is particularly effective [59,60] because it accounts for the electronic effects and energy contributions of water molecules [61–63] that are often crucial in drug binding [58,62] to ion channels [61]. This methodology had been originally validated [58] on a diverse set of 43 target-ligand complexes. In the present study, validation was extended to the available M2 and EP complexes with known activity, and a strong correlation (R^2^ = 0.88, RMSE=0.73 kcal/mol, Fig. 3A) was observed between the QMH-L-calculated ΔG_b_ values and experimental pEC_50_ (logarithm of the 50% effective concentration).

**Fig. 3.**
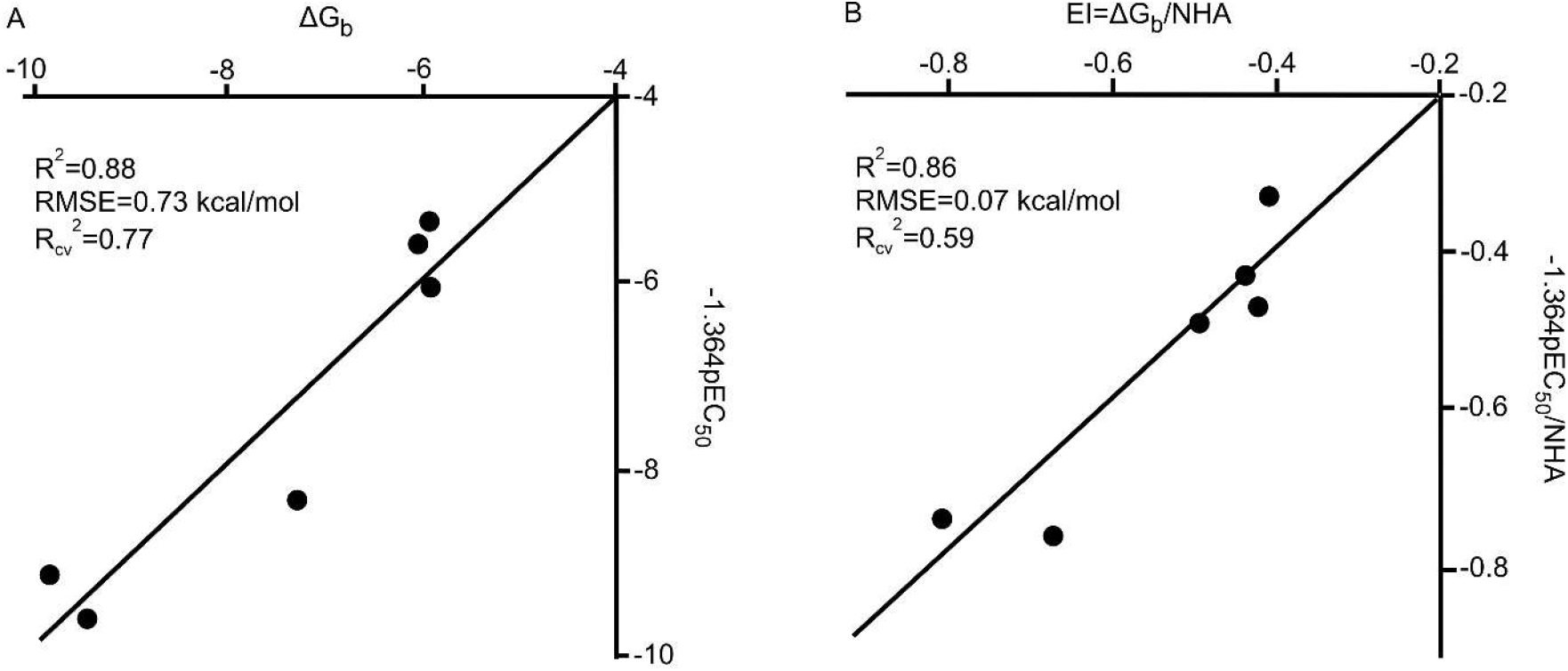
The validation of the QMH-L binding affinity calculations based on complex structures of AMS compounds and ion channels M2 and EP generated by HydroDock in the previous [25], and the present study (Methods). **A**) The correlation plot of experimental antiviral activity (pEC_50_) and calculated binding affinity (ΔG_b_, kcal/mol) is based on experimental EC_50_ data (Table S2) from the literature and QMH-L calculated values (Table S1) of the present study. **B**) The correlation plot was also produced for the calculated EI (kcal/mol) values. The coefficient -1.364 comes from converting pEC_50_ to ΔG_b_ using the equation ΔG_b_ =RT lnEC_50_, where R is the gas constant and T = 298.15 K is the temperature.

Similarly, a strong correlation (Fig. 3B) was observed for the calculated efficiency index (EI = ΔG_b_/NHA, where NHA is the number of heavy atoms [64] in the AMS compound) values. EI is a key metric recommended for scaffold design [64]. Experimental EI was calculated from pEC_50_ (instead of ΔG_b_ in the numerator) and reflects per-atom binding strength (activity) [65]. Along with previous validations [25,58] the results in Fig. 3 show that, beyond their general applicability, the computational tools HydroDock and QMH-L are reliable for designing antivirals, particularly targeting ion channels such as EP.

A co-analysis of the ΔG_b_ values with the corresponding experimental and/or HydroDock-based structures showed that the binding of AMS compounds is weaker to EP than to M2 (Fig. S2). Since the hydrophobic band (HB1, Fig. 2) of EP is wider than that of M2, small molecules like AMA or RIM, which fit perfectly to M2, fail to fill the larger space in EP, resulting in a weaker binding (Table S1) and lower antiviral activity against SARS-CoV-2 (Table S2). Among the tested AMS compounds, arterolane (ART), a repositioned anti-malaria[66] drug has the most negative ΔG_b_ and the strongest binding to EP (Fig. 2, Table S1) because its larger scaffold spans both HB1 and HB2 (Fig. 2), leading to the best antiviral activity (Table S2, Fig. 2). However, the amide and amino groups in ART are also positioned in HB1 (Fig. 2), which makes having these hydrophilic groups in this hydrophobic environment energetically unfavorable. HydroDock calculations (Fig. 2) also showed that three water molecules occupying the HB1 space around the amino group stabilize ART’s binding to EP, even in this unfavorable environment.

Based on the SAR results above from Step 1, we concluded that extending the AMS along the ion channel’s vertical axis improves binding strength and antiviral activity only if the extension isn’t just filling empty cavities in EP. The extension only makes sense if the AMS interacts with consecutive hydrophobic and hydrophilic bands (as ART in Fig. 2) and water molecules inside EP.

### 2 Virology and cytology laboratory verification setup

As a further preparatory step for verifying the new AMS compounds (Steps 3 and 4), virology and cytology laboratory experiments (Fig. 4) were set up and tested with the known drugs listed in Fig. 1 (grey). Robust colorimetric assays [67,68] were used to experimentally verify the antiviral activity (blocking of *in vitro* viral replication) of all AMS (Fig. 1) predicted by the computational design. The antiviral activities of AMS were expressed as the 50% effective concentration (EC_50_), a widely used measure in the literature [69–72]. The assays can readily measure the inhibition of viral cytopathic effect by AMS against various SARS-CoV-2 variants, as shown for ART (Fig. 4A). The effect of ART is also evident in a series of microscopic images of Vero E6 cell cultures (Fig. 4B) used for the EC_50_ measurements, where a considerable difference was observed among the baseline viability of healthy cells, the AMS-treated cells, and the infected cells. The Vero E6 cell line was also used to determine the 50% cytotoxic concentration (CC_50_), which is required to calculate the safety index (SI = CC_50_/EC_50_) for AMS. In some cases, SI< 1 values occur for the delta and omicron variants. Since the sequence of the transmembrane region of EP among different variants is identical (Fig. 2B), these SI<1 values are not due to different binding (ΔG_b_) of the AMS to EP across various variants. (It is likely because of the different replication kinetics of the variants [73]; for example, delta was found to be dominant over alpha [73], which is reflected in our results, where SIs for delta are generally lower compared to alpha for the original four AMS in Table S3.)

**Fig. 4.**
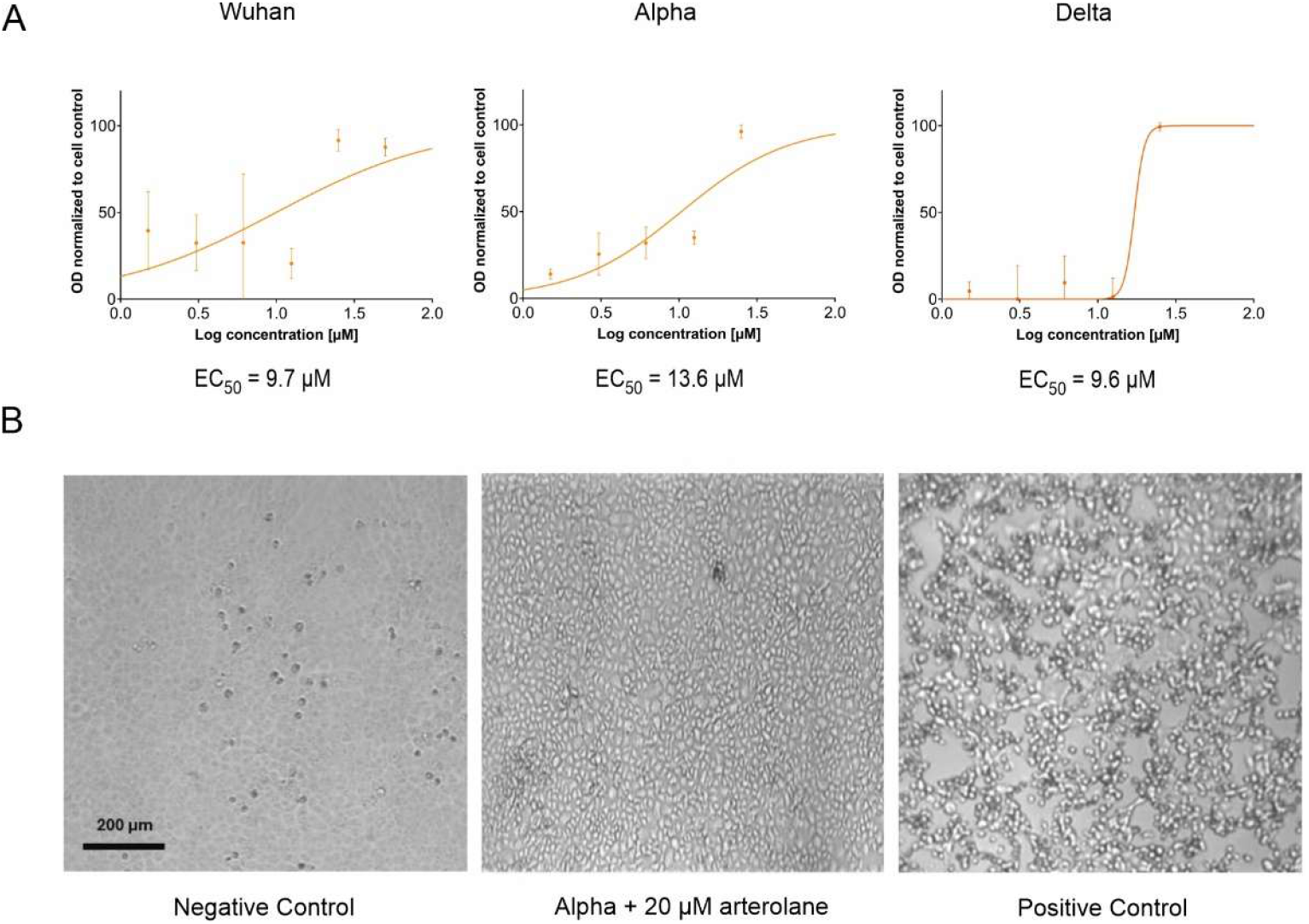
ART inhibits SARS-CoV-2 replication in vitro. A) Wuhan, alpha, and delta strain-infected (labeled accordingly) cells were treated with ART. A sigmoid curve is fitted to the measured optical density data. The standard error of measurement of three replicates is shown with error bars. **B**) Microscopy images show antiviral activity. The microscopy images illustrate the cytotoxic effects of viral infection and their reversal by arterolane treatment. Granule-like appearances indicate cell death. No cytopathic effects were observed after 3 days of Omicron infection with ART treatment, even at the highest concentration (20 µM), where ART showed no cytotoxicity.

The laboratory measurements verified that all four AMS drugs (grey in Fig. 1) have an SI > 1 for the alpha variant (Table S3) as expected based on previous experiments [69,70,72]. Regarding cytotoxicity, AMA is the least toxic (highest CC_50_, Table S3), while ART is the most toxic among the four drugs (Fig. 1B) used as references in this study. The CC_50_ values for the four reference drugs closely align with data from previous research (R^2^=0.97, Table S2, Fig. S3). Overall, our virology and cytology laboratory experiments are consistent with previous laboratory results for the reference drugs and provide a reliable testing platform for the new compounds introduced in the next steps.

### 3 Extension of the AMA scaffold

Building on the previous SAR findings (Step 1), the AMS was redesigned to reduce issues related to the chemical synthesis, accessibility and cost of spiro-compounds like ART and spiro-adamantyl amine [25] (SPA, Fig. S4), while also improving the binding of the base compound AMA to EP. (At the time of preparing this study, ART was purchased at a very high price, and SPA was not available on the market.) To achieve this, the adamantyl ring of AMA was extended with a small linker and a (substituted) phenyl group (Fig. 5, Fig. S4) instead of the spiro-system. This redesign produced two new AMS with an extended scaffold: N-phenyladamantane-1-carboxamide (PAC, Fig. 1B) and 4-(adamantan-1-ylethynyl)aniline (ETA, Fig. 1B), each featuring a different linker (amide and ethynyl, respectively).

**Fig. 5.**
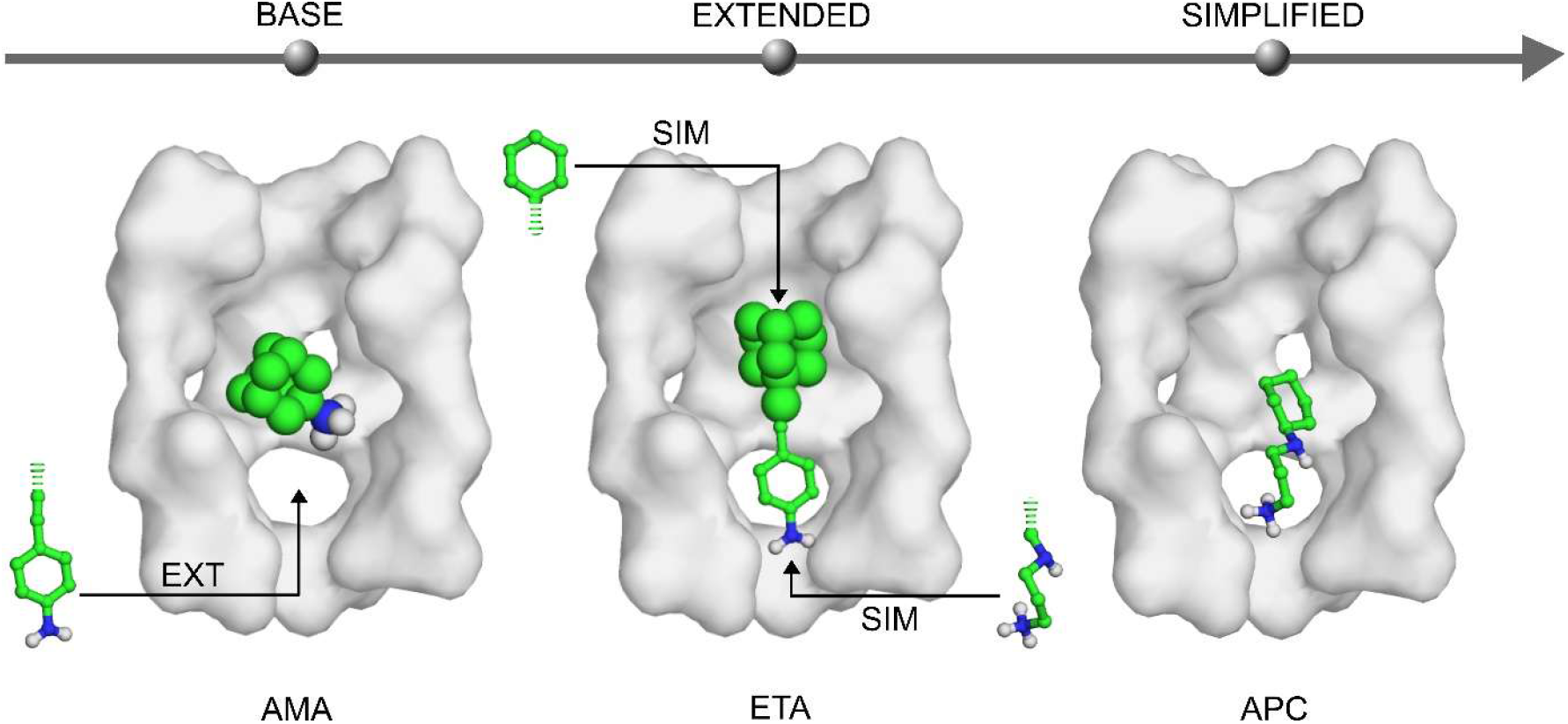
The scaffold redesign strategy used in this study, showing the transformations from AMA to ETA to APC in the AMS series. Water molecules are omitted for clarity, but they are included in the detailed figures. The adamantyl group of AMA is depicted in a space-filling model, and the new groups added during the redesign are shown in ball-and-stick models.

HydroDock modeling shows that these rigid, linear molecules are perfectly sized to span both the HB1 and HB2 regions of EP (Fig. 6). The geometric design eliminated the problematic contacts (Fig. 2) involving the amide and amino groups of ART. While PAC also contains an amide group in the middle, it interacts with the hydrophilic band and a water molecule (Fig. 6A). The amino group of ETA binds to another water molecule within the hydrophilic band of EP, above HB 1. The phenyl groups of PAC and ETA bind to L18 and L23 in the small HB1 (Fig. 6). The bulky adamantyl groups bind to V25 in the larger HB2 (Figs. 2 and 6), similar to ART (Fig. 2). The good structural fit of PAC and ETA to EP is also reflected in an improved ΔG_b_ relative to AMA, RIM, and MEM. While PAC and ETA are considerably smaller than ART, their EI values are comparable, suggesting that a simplified, well-aligned scaffold can outperform larger, more complex molecules.

**Fig. 6.**
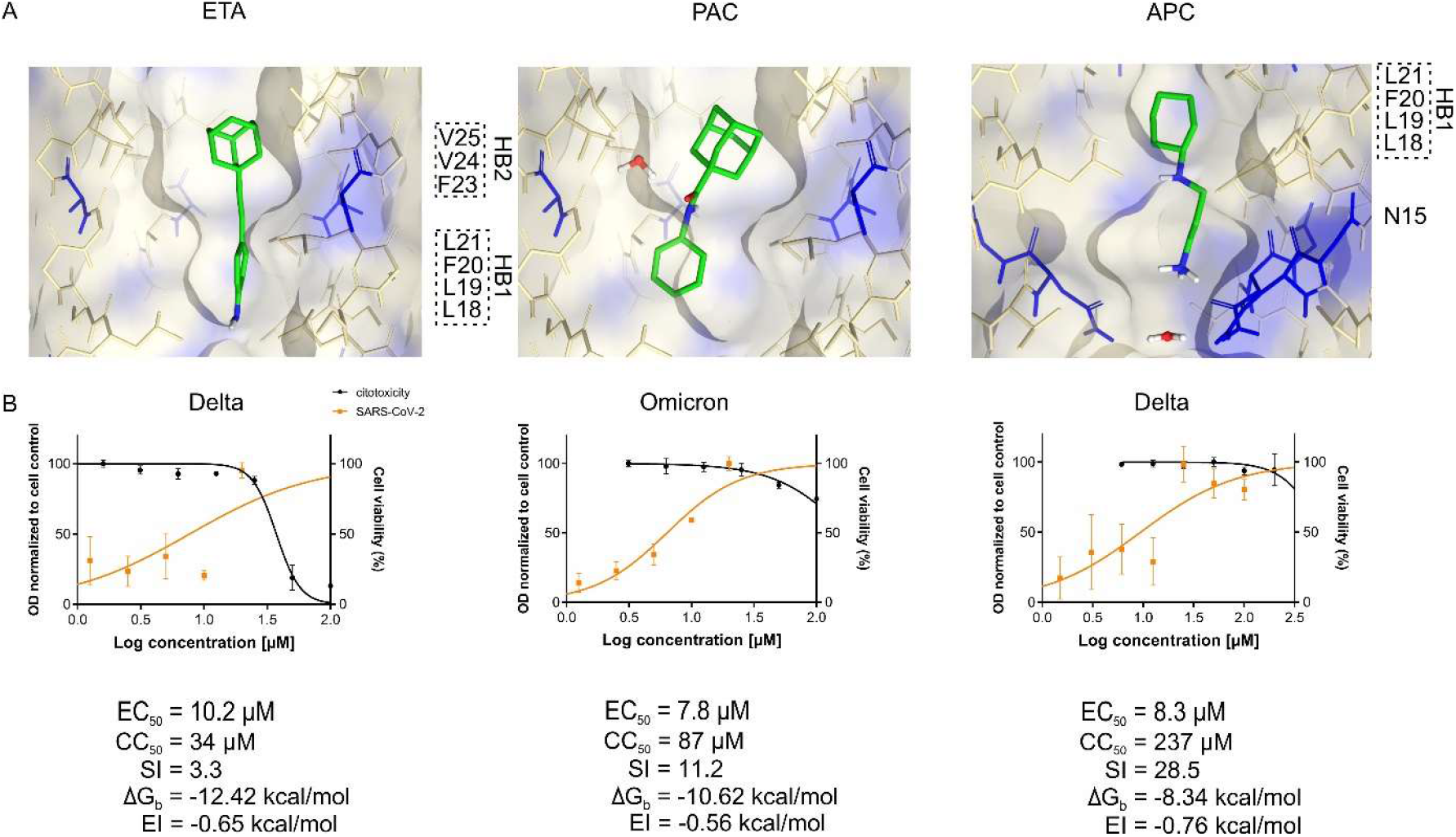
Top results of the simplified scaffold redesign strategy from this study. A) The binding modes of ETA, PAC, and APC to EP as produced by HydroDock. EP is shown as a surface, while ETA, PAC, and APC are represented as green sticks; water molecules are depicted as white and red balls and sticks. B) The results from virology and cytology laboratory tests of ETA, PAC, and APC. The orange curves indicate the inhibitory effects on SARS-CoV-2 replication, and the black curves show the cytotoxicity measurements. The bottom of the figure points toward the extraviral space.

**Figure 7:**
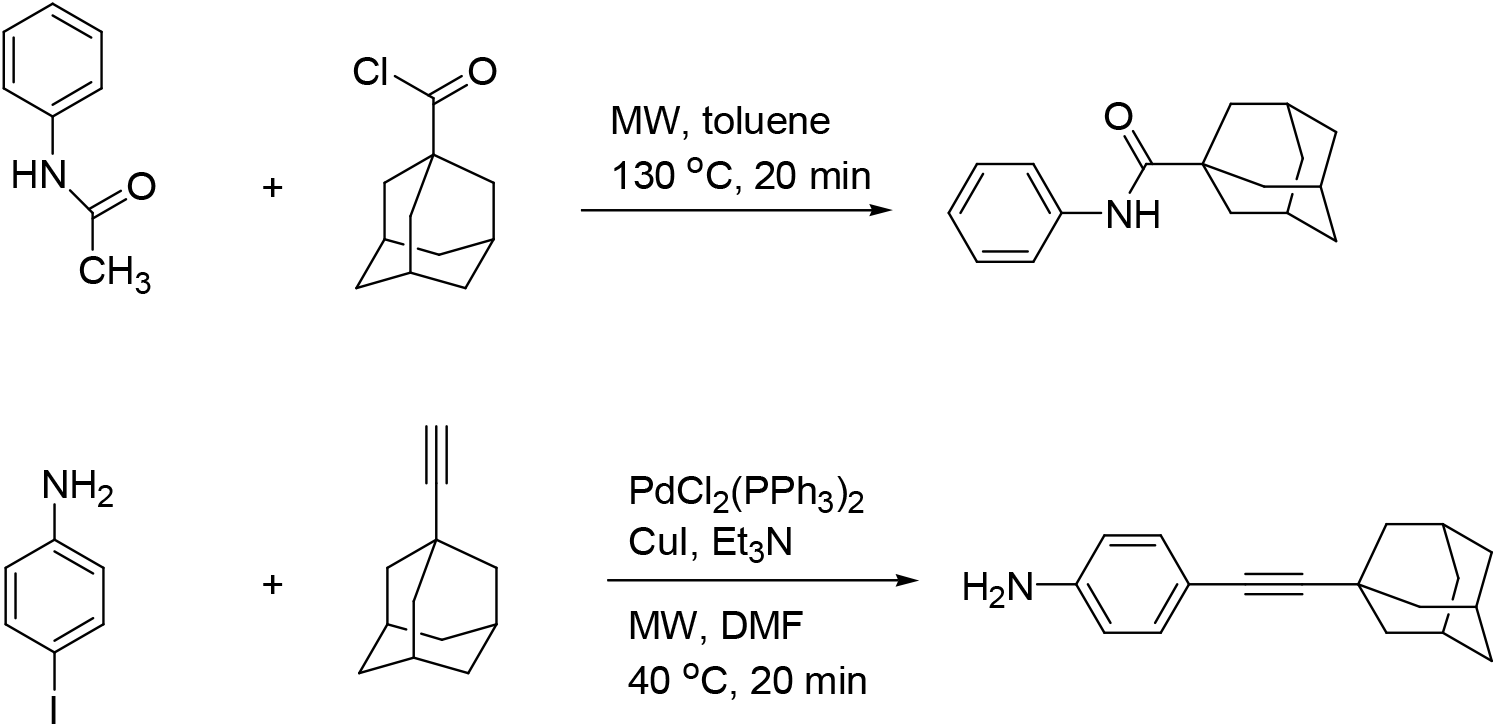
Synthesis of **PAC** (top) and **ETA** (bottom)

The experimental antiviral activities of PAC and ETA were tested against SARS-CoV-2 for the first time in this study, and the results are shown in Fig. 6. Earlier, PAC was synthesized and tested against the influenza A virus [74]. ETA was previously synthesized as a building block for a series of hypoxia-inducible factor 1 inhibitors [75]. We found that ETA exhibits higher antiviral activity against the delta variant than AMA, RIM, and MEM (Table S3). PAC has better SI values against the delta, omicron, and Wuhan variants than those of all previously studied compounds. Building on SAR insights (Step 1), the scaffold extension (Step 3) improved activity and SI for the new AMS compared to AMA. However, the high synthetic cost of the advanced C(sp^3^)–H functionalization [57] of the adamantyl precursors still limits their usefulness.

### 4 Simplification of the scaffold

To enhance synthetic accessibility and maintain (or improve) SI and antiviral activity, the adamantyl scaffold was simplified to either a phenyl or a cyclohexyl group (Fig. 5).

The 4-tert-butyl-cyclohexylamine (BCA, Fig. 1), and trans-4-bocamino cyclohexylamine (BOC, Fig. 1). 3-methoxytyramine (MOX, Fig. 1) is assumed based on the use of a phenyl ring. Between the hydrophobic moiety and the amino group, there are two atoms in MOX, and no atoms in BCA and BOC. The antiviral potential of these compounds against SARS-CoV-2 is reported here for the first time. The laboratory experiments demonstrate that BCA is highly cytotoxic; therefore, it was not tested for antiviral activity. MOX and BOC are not cytotoxic; however, they are ineffective against SARS-CoV-2 *in vitro* (as expected, since their binding modes do not align with the hydrophobic/hydrophilic regions of EP).

The simplification of the adamantly scaffold to a cyclohexyl group yielded N-3-aminopropyl cyclohexylamine (APC). The binding mode of APC to EP was calculated using HydroDock (Fig. 6). In the four-atom linker between the cyclohexane ring and the terminal amino group of APC, there is an imino group (Fig. 1B), which can interact with the backbone carboxyl of N15 below the HB1 of EP (Fig. 6). The three-carbon alkyl linker of APC provides sufficient space for the amino group to interact with the N15 side chain and a water molecule below HB1 (Fig. 6). The cyclohexyl ring interacts with the side chains of L18 and L21 in HB1, similar to the adamantane scaffold of the base compound AMA (Fig. 5). Consequently, the corresponding parts of APC align well with the hydrophilic and hydrophobic regions of EP, similar to PAC or ETA (Step 3). The calculated ΔG_b_ of APC is more favorable than that of the three drugs AMA, RIM, and MEM (Table S1) and is comparable to PAC. Additionally, the EI of APC is the best among all compounds (Table S1, Fig. 6), indicating that APC is a promising candidate for further laboratory testing.

In a previous study, APC was tested against malaria [76], but not against viruses. Our laboratory results confirmed that APC has low cytotoxicity (Table S3, Fig. 6) and is highly effective against SARS-CoV-2 *in vitro* (Table S3). Among all tested compounds, APC shows the most potent antiviral activity against the delta strain (Table S3) and exhibits antiviral activity comparable to ART against the omicron and Wuhan strains. The antiviral activity of APC against the omicron strain is slightly lower than that of PAC but still surpasses AMA, RIM, and MEM. The SI of APC exceeds 22 and 28 against the Wuhan and delta strains, respectively, and is over two against the alpha and omicron strains. Since APC is a much simpler compound than others with adamantyl-based AMS (Fig. 1), it provides a synthetically more feasible and still active scaffold for the future design of broad-spectrum antivirals targeting ion channels.

## 5 Conclusions

Common high-throughput screening [3] and drug repositioning have many limitations in antiviral drug research [14,77]. Instead, in the present study, we redesigned the antiviral scaffold through a 4-step target-based engineering approach. In Step 1, SARs were derived using advanced computational methods, HydroDock [25] and QMH-L [58] that can handle water molecules [25,78] and electronic effects for the precise prediction of binding affinity. As a laboratory test system (Step 2), we targeted multiple SARS-CoV-2 variants that encode a non-mutagenic ion channel (EP) with strong potential as a drug target. Based on the SARs we designed an extended AMS (Step 3), resulting in an improved fit within the ion channel and the identification of two new compounds. Finally, the simplification (Step 4) yielded a cyclohexylamine-based minimalist scaffold with good synthetic accessibility. Significant improvements were achieved over approved drugs regarding efficiency and safety. Since the geometry and alternating hydrophobic pattern are standard features of ion channels in various viruses, the final, simplified scaffold will serve as an excellent starting point for broad-spectrum antiviral drug design.

## Materials and Methods

### Molecular engineering

#### Calculation of ligand binding modes

The HydroDock [25] protocol was applied to produce the hydrated target-ligand complexes of AMS bound EP. The M2 and EP bound AMA and RIM structures were used from our previous study [25]. The atomic coordinates EP were acquired from the Protein Data Bank [79] under the accession code 7k3g [28]. The N- and C-terminal ends of the protein were capped in Maestro [80]. Polar hydrogen atoms and Gasteiger-Marsili partial charges were added in AutoDockTools [81]. MEM, ART, PAC, ETA, MOX, APC, BOC, and BCA were built in Maestro, and energy minimized with OpenBabel [82]. The parameter files of the compounds for the molecular dynamics simulations were prepared using the CHARMM-GUI [83].

Dry docking of the compounds to EP was performed in AutoDock 4.2.6. 100 blind docking runs were performed, and the docking box covered the entire surface of the protein. Flexibility was allowed on the ligand side; the Lamarckian genetic algorithm was used. The ligand binding modes were clustered and ranked based on their calculated free energy of binding values.

The hydrated target files were used from our previous study [25]. In the third step of the HydroDock protocol, the 1^st^-ranked, dry-docked binding modes were merged with the hydrated target files. If the 1^st^-ranked, dry-docked binding mode was outside the channel, then the 1^st^-ranked binding mode was used, which is inside the channel.

#### Minimization of the complex

Overlapping water molecules with the ligands were removed in MobyWat [84]. The resulting hydrated target – dry docked ligand complex structures were subject to a five-step energy minimization [25]. The system was solvated in explicit TIP3P water molecules, and counterions were added to achieve neutrality. First, a steepest descent and a conjugate gradient minimization were carried out, followed by a 100-ps molecular dynamics (MD) simulation and a second steepest descent and conjugate gradient minimization, exactly as in [25]. The resulting systems were ready for molecular dynamics (MD) simulations lasting up to 100ns. AMBER [85] force field was used during MD simulations. The temperature was coupled to a constant of 303.15K. The pressure was coupled to a constant of 1 bar. Particle Mesh-Ewald summation was used for long-range electrostatics. Van der Waals and Coulomb interactions were cut off at 11 Å. Position restraints were applied to the Cα atoms of the target protein. The final trajectory containing all atomic coordinates of all frames was converted to a portable DXR binary file. The representative structure of each simulation was selected as the closest match to the statistical average of all ligand frames, as in [25]. There was one exception, in the case of APC, where the frame with the most favorable calculated interaction energy (E_inter_, see next Section) was selected. The binding mode of the energetically most favorable and the representative frames was similar.

#### E_inter_ energy calculation

The energy of the representative binding modes was calculated, including water molecules within 3.5 Å distance from the ligand, and these water molecules were treated as part of the target. Lennard-Jones and Coulomb (Mehler-Solmajer [86]) interaction energies were calculated as in eqs. 1 and 2. The AMBER parameter files were used from the MD simulations, besides the atomic coordinates as input files for energy calculation.

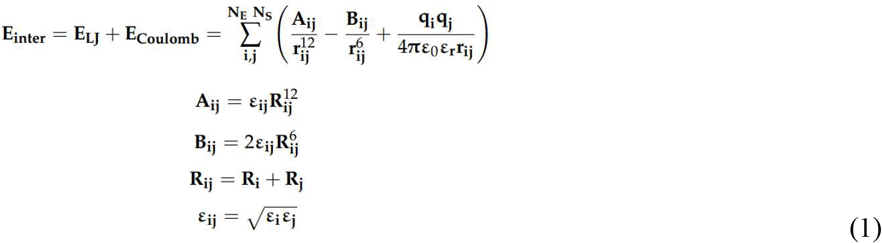

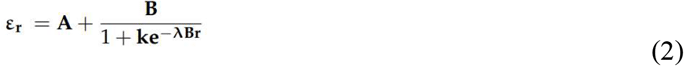

The E_inter_ is the sum of Lennard-Jones (LJ) and Coulomb (Cb) intermolecular interaction energies (1). The Coulomb term was calculated with a distance-dependent dielectric function (2) In (1), ε_ij_ is the potential well depth at equilibrium between the i^th^ (ligand) and j^th^ (protein) atoms; ε_0_ is the permittivity of vacuum; ε_r_ is the relative permittivity (2); R_ij_ is the inter-nuclear distance at equilibrium between i^th^ (ligand) and j^th^ (protein) atoms; q is the partial charge of an atom; rij is the actual distance between the i^th^ (ligand) and j^th^ (protein) atoms; NE is the number of protein atoms; NS is the number of ligand atoms. In (2), B = ε_0_ -A, ε_0_ is the dielectric constant of water at 25°C, and A, λ, and k are parameters.

#### ΔG_b_ calculation

For the representative frames, a quantum mechanics-based change in free energy of binding (ΔG_b_) was calculated as described in [58]. Briefly, the water molecules within 3.5 Å distance from both target and ligand were kept, and the heat of formation (ΔfH) of the hydrated target-ligand complex, the dry target, and the dry ligand were calculated using MOPAC [87] quantum chemistry software. According to Hess’ law (eq. 3), the ΔfH of the target, the ligand, and the water molecules (-65.2 kcal/mol multiplied by the number of water molecules) were subtracted from the ΔfH of the complex, resulting in a calculated heat of reaction (ΔrH) as described in [58]. Then, the number of heavy atoms was divided by the number of atoms to result in the NHA/NA descriptor necessary for the calculation of ΔG_b_ according to (eq. 4, [58]). The calculated ΔG_b_ values are shown in Table S1.

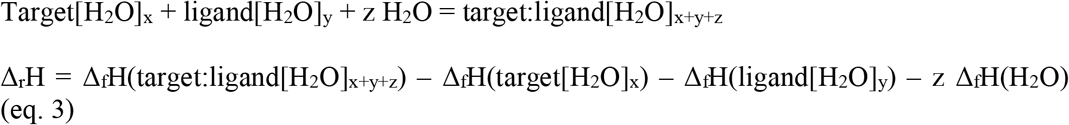

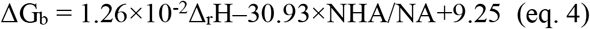

### Chemical syntheses

While the synthesis of PAC and ETA has been previously documented in the literature [74,75], we also achieved a significant reduction in reaction times by using one-pot syntheses and microwave-assisted conditions (Sonogashira coupling yielded a notable increase in ETA yield), in line with green chemistry principles. Thus, PAC and ETA are simpler extensions of AMA, providing an easily synthesizable scaffold for AMS alternative to the expensive, complicated ART.

#### N-Phenyladamantane-1-carboxamide (PAC)

1-Adamantanecarbonyl chloride (199 mg, 1.00 mmol), acetanilide (135 mg, 1.00 mmol), and toluene (3 mL) were added to a 10 mL microwave vessel under a nitrogen atmosphere. The mixture was heated in a microwave reactor at 130 °C for 20 min under stirring. After cooling to rt, the solvent was evaporated *in vacuo*, and the residue was purified by flash chromatography with 20% ethyl acetate/80% hexane as eluent. Compound **PAC** was identical to the compound described in the literature [74].

#### 4-(adamantan-1-ylethynyl)aniline (ETA)

4-Iodoaniline (219 mg, 1.00 mmol), PdCl_2_(PPh_3_)_2_ (70 mg, 0.1 mmol), CuI (19 mg, 0.1 mmol) and DMF (3 ml) were added under nitrogen atmosphere, then Et_3_N (0.84 ml, 6 mmol) was added and the mixture was stirred at 40 °C for 10 min in a 10 mL microwave vessel. 1-Ethynyladamantane (160 mg, 1.00 mmol) was added, and the mixture was heated in a microwave reactor at 40 °C for 30 min under stirring. The solvent was evaporated in vacuo. The residue was purified by flash chromatography with 20% ethyl acetate/80% hexane as eluent. Compound **ETA** was identical to the compound described in the literature [75].

### Chemical analyses

Thin-layer chromatography was performed on silica gel 60 F254 (layer thickness 0.2 mm, Merck). The spots were detected with I_2_ or UV (365 nm). Flash chromatography was performed on silica gel 60, 40–63 μm (Merck). Reactions under microwave irradiation were carried out in the Anton Paar Monowave 400 microwave reactor. ^1^H NMR spectra were recorded in DMSO-d_6_ solution with a Bruker DRX-500 instrument at 500 MHz. ^13^C NMR spectra (Figures S5 and S6) were recorded with the same instrument at 125 MHz under the same conditions.

#### PAC

Compound **PAC** was obtained as a white solid (232 mg, 91%). Anal calcd. for C_17_H_21_NO: C, 79.96; H, 8.29. Found: C, 80.05; H, 8.34. Mr: 255.35. ^1^H NMR (500 MHz, DMSO-d_6_) δ ppm:1.71(broad s, 6H), 1.90(overlapping signals, 6H), 2.02(broad s, 3H), 7.02(t, 1H, *J* = 7.4 Hz), 7.27(t, 2H, *J* = 7.4 Hz), 7.64(d, 2H, *J* = 7.4 Hz), 9.07(s, 1H, NH); ^13^C NMR (125 MHz, DMSO-d_6_) δ ppm: 27.6 (3C, 3xCH), 35.9 (3C, 3xCH_2_), 38.2 (3C, 3xCH_2_), 40.8 (C), 120.1 (2C, 2xCH), 123.0 (CH), 128.2 (2C, 2xCH), 139.2 (C), 175.8 (C).

#### ETA

Compound **ETA** was obtained as a white solid (224 mg, 89%). Anal calcd. for C_18_H_21_N: C, 86.01; H, 8.42. Found: C, 86.09; H, 8.47. Mr: 251.37. ^1^H NMR (500 MHz, DMSO-d_6_) δ ppm:1.67(s, 6H), 1.86(overlapping signals, 6H), 1.94(s, 3H), 5.32 (s, 2H, NH_2_), 6.46(d, 2H, *J* = 7.4 Hz), 6.97(d, 2H, *J* = 7.4 Hz); ^13^C NMR (DMSO-d_6_) δ ppm: 27.3 (3C, 3xCH), 29.4 (C), 35.7 (3C, 3xCH_2_), 42.7 (3C, 3xCH_2_), 80.2 and 94.6 (C≡CH), 109.4 (C), 113.4 (2C, 2xCH), 132.1 (2C, 2xCH), 148.3 (C).

### Virology and cytology laboratory measurements

#### Cell lines and virus strains

Vero E6 cells (African Green Monkey renal epithelial cells; ATCC cat. no. CRL-1586) were maintained in Dulbecco’s Modified Eagle Medium (DMEM) (Lonza, Basel, Switzerland) supplemented with 10% heat-inactivated fetal bovine serum (FBS) (Gibco, Waltham, MA, USA) and 1% penicillin–streptomycin (Lonza, Basel, Switzerland) at 37°C and 5% CO_2_ in a humidified atmosphere. The Vero E6 cell line can be infected with SARS-CoV-2 [88] and the study was performed with different variants of SARS-CoV-2: P.2 - mentioned later as wuhan variant; B.1.617 - hereinafter referred as delta variant; B.1.1.529 - mentioned later as omicron variant; B.1.1.7. - referred to as the alpha variant. The experiments were performed in a biosafety level 4 (BSL-4) laboratory.

#### Reagent preparation

Amantadine-hydrochloride (AMA, 97% purity), memantine-hydrochloride (MEM, 98% purity), and 1-(1-adamantyl)ethylamine-hydrochloride, also called rimantadine (RIM, 99% purity, racemic), were purchased from Merck (Darmstadt, Germany). arterolane (ART, 98% purity), 3-methoxytyramine (MOX, 98.36% purity), and N-(3-Aminopropyl)cyclohexylamine (APC, 99,37% purity) were purchased from MedChemExpress (USA). Trans-4-boc-aminocyclohexylamine (BOC, 97% purity) and 4-tert-Butylcyclohexylamine (BCA) were purchased from ThermoFisher (USA). AMA, MEM, RIM, ART, MOX, APC, BOC, BCA, PAC, and ETA were solubilized in sterile water or DMSO, and 10 mM stocks were made of them. The drugs were further diluted with medium to reach working concentrations.

#### Cytotoxicity assay

To determine the cytotoxicity of test drugs, a 3-(4,5-dimethylthiazol-2-yl)-2,5-diphenyltetrazolium bromide (MTT) cell viability assay was performed, and cell cytotoxicity was examined with Cell Proliferation Kit I (Roche, Switzerland). The MTT assay is a quick colorimetric method to assess the viability of cells by measuring their metabolic activity. Viable cells convert MTT to formazan via cellular oxidoreductases, leading to a color change measured by spectrophotometry (absorbance). During the cytotoxicity test, VeroE6 cells were seeded into a 96-well tissue culture plate at a density of 3 × 10^4^ cells per well and were incubated overnight. After that, cells were treated with compounds diluted at the indicated concentrations. RIM, MEM were used at seven different concentrations ranging from 100 µM to 400 µM. AMA was used at six different concentrations ranging from 150 µM to 800 µM. ART, MOX, APC, BOC, BCA, PAC, and ETA were used at fourteen different concentrations ranging from 3 µM to 600 µM. The different concentration values were selected based on the preliminary microscopically observed toxicity values (data not mentioned in the manuscript). After 72 h of treatment, cells were washed with DMEM, and MTT Labeling reagent was added. After 4 h incubation at 37 °C, the Solubilization solution was added and incubated overnight. The absorbance of formazan was measured at 570 nm with Crocodile 5in1 mini Workstation (Berthold, Germany). The 50% cytotoxic concentration (CC_50_) values were calculated using GraphPad Prism version 8.00 software (GraphPad Software, San Diego, CA, USA) using non-linear regression. Three technical replicates of each concentration were used. Three biological replicates of the positive and negative controls were used. The selectivity index (SI) was calculated by dividing the EC_50_ (calculated as described below) by the CC_50_.

The use of cytotoxicity tests was important to find out the concentration of the drugs that do not cause cell death, but their concentration is high enough to inhibit virus replication.

#### Cytopathic effect inhibition assay

Antiviral assays were performed under BSL-4 conditions, and viral stocks were prepared as previously described [88]

To test the effect of drugs on virus replication, Vero E6 cells were seeded into 96-well plates at a density of 3 × 10^4^ cells per well the day before the antiviral experiment. Cells were treated with the test compounds at different concentrations (previously described). Immediately after treatment, cells were infected with different variants of SARS-CoV-2 at the multiplicity of infection (MOI): 0.5. MOI is the ratio of agents to infection targets. Cells were incubated for 30 min at 37 °C, then the supernatant was replaced with fresh maintenance media (DMEM, 2% FBS, 1% Pen/Strep) supplemented with the compounds at the appropriate concentration. After 72 h of treatment, virus-induced cytopathic effects (CPE) were microscopically examined (data not mentioned in the manuscript), cells were washed with DMEM, and MTT Labeling reagent was added. After 4 h incubation at 37 °C, the Solubilization solution was added and incubated overnight. The absorbance of formazan was measured at 570 nm with Crocodile 5in1 mini Workstation (Berthold, Germany). The 50% effective concentration (EC_50_) was calculated using GraphPad Prism version 8.00 software (GraphPad Software, San Diego, CA, USA) for non-linear regression. Three biological replicates of each concentration were used. Treatment plates included biological replicates of the positive and negative controls; infected/nontreated and noninfected/nontreated controls were used.

#### Microscopy

Vero E6 cells were seeded on 96 96-well plate, infected with variants, as alpha, omicron, and delta variants at an MOI of 0.5, and treated with ART in parallel with infection. The images (**Fig. 4**) were taken with Omni™, a live-cell analysis platform.

#### Statistical analysis

Sigmoidal concentration-response curves were fitted, EC50 values and plots were generated using GraphPad Prism version 8.00 software (GraphPad Software, San Diego, CA, USA).

## Supporting Information

Supporting Information: ^1^H NMR spectra for synthesized compounds (DOC).

## Acknowledgments

The manuscript is dedicated to the memory of Prof. Dr. Ferenc Jakab. We acknowledge the Digital Government Development and Project Management Ltd. for awarding us access to the Komondor HPC facility based in Hungary.

## Author contribution

B.Z.Z and C.H. elaborated the strategy, designed theoretical calculations, interpreted the data, and wrote the manuscript. C.H. supervised the project. E.M. synthesized compounds and wrote the corresponding part of the manuscript. F.F, Z.K., K.L., M.M., B.Z., K.A. performed laboratory experiments and wrote the corresponding part of the manuscript.

B.Z.Z and C.H. acquired funding.

## Data availability

The authors declare that all the data supporting the findings of this study are available within the paper and its Supplementary Data, and deposited at: https://zenodo.org/records/18323738.

